# Bacterial sirtuin CobB and PRPP synthase crosstalk in the regulation of protein acetylation in *Escherichia coli*

**DOI:** 10.1101/2020.12.09.417477

**Authors:** Beata M. Walter, Joanna Morcinek-Orłowska, Aneta Szulc, Andrew L. Lovering, James R.J. Haycocks, Manuel Banzhaf, Monika Glinkowska

**Affiliations:** Structural Biology Laboratory, Intercollegiate Faculty of Biotechnology, University of Gdansk and Medical University of Gdansk, 80-307 Gdansk, Poland; Department of Bacterial Molecular Genetics, Faculty of Biology, University of Gdansk, Gdansk, Poland; Institute of Microbiology & Infection and School of Biosciences, University of Birmingham, UK; Newcastle University Biosciences Institute, Faculty of Medical Sciences, Newcastle University, Newcastle upon Tyne, NE2 4HH, UK

**Keywords:** Protein lysine acetylation/ deacetylase/ sirtuins/ NAD metabolism/ phosporibosyl pyrophosphate synthase

## Abstract

In bacteria, post-translational acetylation of lysine residues affects the activities of numerous proteins, thus constituting an important regulatory mechanism that influences bacterial growth and adaptation to various environments. Conversely, the modification level of some lysine residues is controlled by deacetylases that remove the acetyl group. Most bacterial deacetylases are homologous to eukaryotic sirtuins, which utilize NAD+ as a cofactor. The *Escherichia coli* genome encodes a single sirtuin – CobB. However, both the regulation of CobB activity and the role that reversible acetylation plays in the control of protein functions and cellular pathways are not fully understood.

In this study, we demonstrate that CobB forms a stable complex with PRPP synthase Prs. PRPP synthase is responsible for providing precursor metabolite for the synthesis of certain amino acids as well as nucleotides, including the cofactor NAD+. This association stimulates the deacetylation rate by CobB and protects it from inhibition by its reaction byproduct nicotinamide. We present evidence indicating that acetylation of Prs is physiologically significant and impacts *E. coli* metabolism and global protein acetylation level, although it does not affect CobB binding *in vitro*. In turn, Prs can be deacetylated by CobB, while formation of CobB-Prs complex formation safeguards PRPP synthase activity under non-optimal conditions. We propose that formation of the complex regulates deacetylation of other protein substrates by CobB.

## Introduction

Lysine acetylation is a post-translational protein modification regulating a wide range of protein functions. It has been extensively studied in eukaryotic cells and has recently been found to be important for protein regulation in prokaryotes as well (1, 2). In bacteria, the level of protein acetylation is a result of two opposing processes. Proteins can be enzymatically acetylated by lysine transacetylases or non-enzymatically acetylated by acetyl phosphate (AcP), respectively (3–5). On the other hand, acetyl groups can be removed from acetyl-lysine residues (AcK) by deacetylases, which are mostly NAD+-dependent homologs of eukaryotic sirtuins (6, 7).

The *E. coli* CobB deacetylase can remove acetyl groups resulting from both enzymatic and non-enzymatic acetylation(8). However, *in vivo* most AcP-mediated acetylation modifications are irreversible(4, 5, 8). Despite this, at least 69 lysine residues of 51 proteins were found to be significantly more acetylated in a strain lacking *cobB* compared to the wild type. It has been suggested that CobB activity primarily affects low stoichiometry sites, similarly to human Sirt3 activity (9, 10). This implies that CobB-mediated deacetylation operates on only a small fraction of given protein molecules and could play a role in regulating specific subsets of proteins spatially or temporally distinct from the rest(11, 12).

CobB is a highly conserved protein lysine deacetylase among bacteria (13, 14). It utilizes NAD+ to remove the acetyl group from AcK residues and produces nicotinamide (NAM) and 2”-O-acetyl-ADP-ribose as byproducts. CobB is also the only sirtuin-like deacetylase identified in *E. coli* (15). Its activity affects many cellular functions (4, 5, 10, 16, 17) including gene expression (18–20), cell cycle (21), metabolism (17, 22, 23), stress responses (3, 17, 24), motility and pathogenicity (25). Additionally, CobB can also act as desuccinylase, de-2-hydroxyisobutyrylase and lipoamidase (26–28).

Despite its important role, factors affecting the rate of protein deacetylation or selection of regulated lysine positions in bacteria are poorly understood. The levels of NAM – one of the products released after acetyl group removal by sirtuins (29), as well as NADH and c-di-GMP (30). In eukaryotes, interplay between sirtuins activity and metabolism was established, showing crucial regulatory role for sirtuins substrate NAD+ in controlling chromatin structure, DNA repair, lifespan and circadian rhythm (31, 32). In bacteria, such interplay seems less obvious. Bacterial sirtuins on one hand hand exhibit low K_m_’s for NAD+ whereas its intracellular concentrations may reach up to the mM range (33). This may jointly result in lower sensitivity of bacterial sirtuins to regulation by NAD+ than their eukaryotic homologues.

To shed light on the regulation of protein deacetylation in *E. coli*, as well as its coordination with other cellular processes, we investigated the CobB deacetylase protein interaction network by affinity purification coupled with mass spectrometry (AP-MS). We further characterized the most prominent interaction between CobB and the phosporibosyl pyrophosphate (PRPP) synthase Prs, whose primary activity fuels metabolic pathways with nucleotide precursors for nucleic acids and cofactor synthesis.

In this work we propose that coordination between protein modification via reversible lysine-acetylation and metabolic state is exerted through crosstalk of *E. coli* the CobB deacetylase with Prs. Namely, we present evidence that CobB forms a stable complex with Prs increasing deacetylase catalytic rate. As result, CobB shows also increased resistance towards inhibition by reaction’s product – NAM. Formation of this complex is not dependent on Prs acetylation status *in vitro,* but we show data strongly suggesting physiological significance connected with modification of Prs by lysine acetylation. Changes to acetylation status of Prs K182, and to a lesser extent also K231, affect *E. coli* growth, global protein acetylation and CobB activity *in vivo*. We also show that, under non-optimal conditions, the rate of PRPP synthesis by Prs may also be influenced by CobB. Based on those observations, we propose that this mutual dependence links nucleotide/NAD+ metabolism and cellular energetics to protein acetylation.

## Results

### CobB interacts with phosphoribosyl pyrophosphate synthase *in vivo* and *in vitro* irrespective of the Prs acetylation status

One way to understand the regulation of CobB-mediated protein deacetylation is to identify its protein interaction partners under physiologically relevant conditions. Some of the potential CobB interactants were identified with the use of a protein microarray consisting of the majority of the *E. coli* proteome (∼4000 proteins, non-acetylated) (34). In this work, we used an approach based on immunoprecipitation pull-down (Fig. 1A), dubbed sequential peptide affinity purification (SPA) (35). In this method, proteins are labeled with a double-affinity tag consisting of triple FLAG epitope and calmodulin binding protein (CBP), separated by a protease cleavage site. After immunoprecipitation from cell lysates with anti-FLAG antibodies, protein complexes are released from the resin by protease digest and re-purified using a calmodulin sepharose column. As a bait, we used chromosomally expressed (from the native promoter), C-terminally SPA-tagged CobB. Subsequently to the pull-down, interaction partners were identified by liquid chromatography coupled to tandem mass spectrometry (LC-MS/MS). The experiments were carried out using cells grown to stationary phase in a rich undefined medium (LB) and minimal medium containing acetate as a carbon source, both conditions supporting high protein acetylation level. Under those conditions, the major component of the purified CobB complex was the phosphoribosyl pyrophosphate synthase Prs (Supplementary Table 1). This interaction has been previously suggested by others in a large-scale study of protein-protein interactions, confirming validity of our results (36).

**Figure 1.**
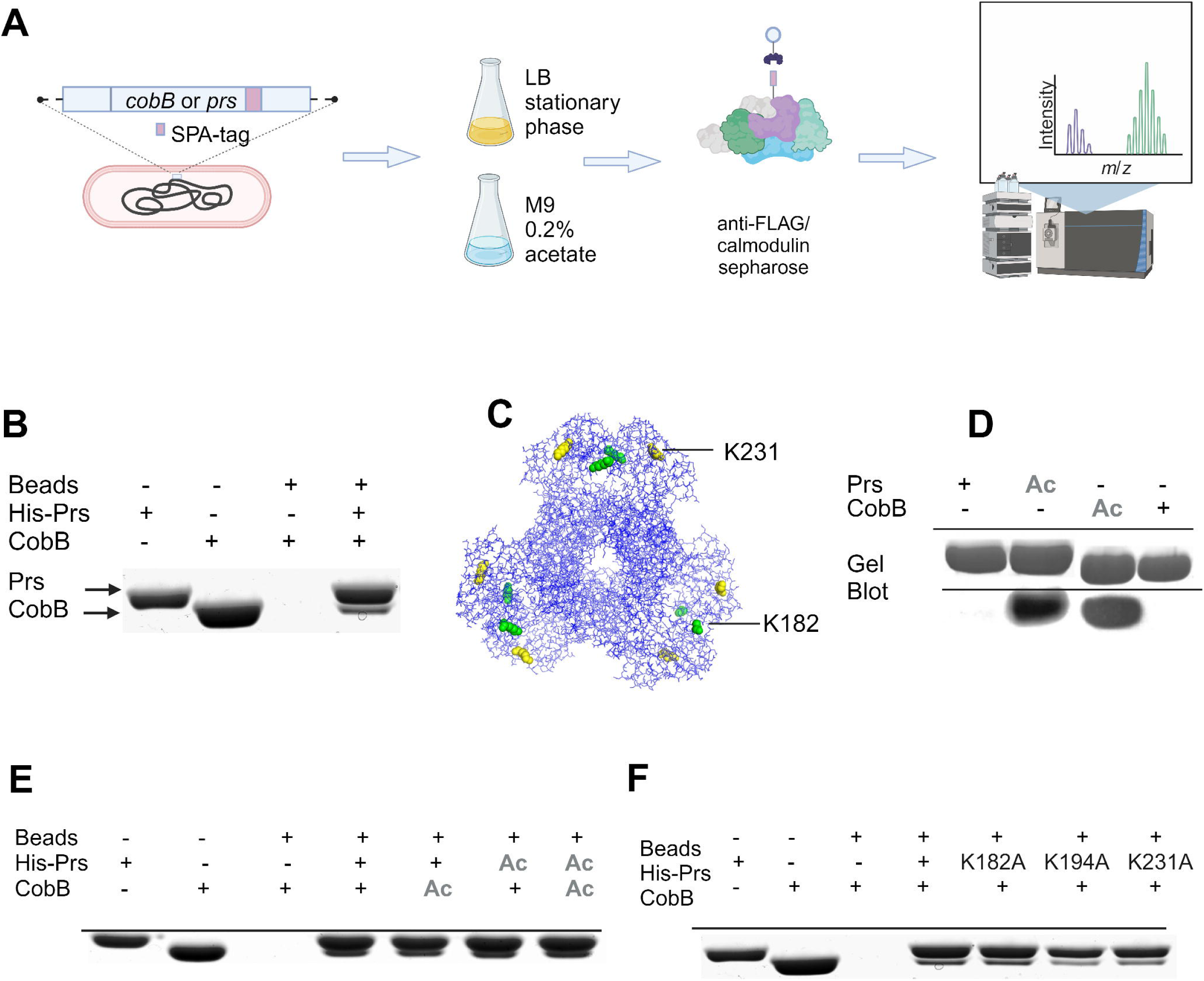
CobB interacts with Prs *in vitro* and in cell lysates irrespective of Prs acetylation status. **(A) CobB and Prs interact in cellular lysates.** Protein-protein interactions were analyzed by AP-MS using SPA-tagged, chromosomally expressed protein variants. **(B) CobB and Prs form a complex *in vitro*.** His_6_-Prs interacts with CobB in a pull-down assay. His_6_-Prs was pre-bound to Ni^2+-^coated magnetic beads. After washing off the excess of Prs, the beads were incubated with CobB. Unbound protein was removed by washing and the beads were resuspended in a loading dye and separated in 10% SDS-PAGE gel. Lane 3 shows the extent of CobB binding to magnetic beads in the absence of the bait. **(C)** Prs structure with highlighted acetylable lysine residues K182 and K231 **(D) Prs and CobB can be acetylated *in vitro* with acetyl phosphate.** Prs and CobB were acetylated in the presence of 20 mM acetyl phosphate and excess of the reagent was dialyzed. Acetylation was confirmed by Western blot with anti-acetyl lysine antibodies. **(E) CobB interacts with both chemically acetylated Prs and (F) Prs variants with key acetylable lysine residues substituted by alanines.** Interaction of acetylated proteins was tested *in vitro* in a pull-down assay, as above. The presence of acetylated protein in a reaction was marked with Ac.

We further compared our data with the interactions previously described in a CobB protein microarray study (34). We found that fatty acids synthesis proteins FabB and FabG are significantly enriched in the presence of CobB, in comparison to control samples analyzed by LC-MS/MS. However, other interactants found in this protein microarray-based investigation were not present or enriched in our data set, suggesting that those interactions are of lower affinity or more transient than the one between CobB and Prs, or do not occur under the chosen experimental conditions (Supplementary Table 1).

We confirmed the interaction between Prs and CobB performing the AP-MS experiment in a reversed set-up, where SPA-tagged Prs was used as a bait (Supplementary Table 1). Consistently, we also observed binding of His-tagged Prs (bait) and CobB (prey) in a pull-down experiment consisting of purified proteins (Fig. 1B).

Overall, these results indicate that CobB forms a stable complex with Prs and that the interaction with Prs is likely one of the most prominent protein-protein interactions formed by CobB in *E. coli* cells.

Prs has been previously reported as one of the proteins acetylated *in vivo* at widely conserved positions K182, K194, K231(5, 10, 17) (Fig. 1C; Supplementary Fig. 1). Therefore, we investigated whether Prs interaction with the CobB is affected by acetylation of its lysine residues, particularly the surface exposed K182 and K231. To achieve this, we first repeated the pull-down experiment using acetylated His-tagged Prs and CobB.

Prs was acetylated *in vitro* by reacting with acetyl phosphate, as described before by others (5, 20). Subsequently, successful modification of lysine residues was confirmed by western blot (Fig. 1D). In addition, we verified by mass spectrometry that the lysine residues of Prs that were acetylated are the same ones shown to be modified in this manner *in vivo* (K182, K194, K231)(Table 1). Non-enzymatically acetylated Prs interacted with CobB in a manner indistinguishable from the unmodified protein (Fig. 1E). It is worth noting that the pull-down reactions did not contain NAD+, disabling deacetylation of Prs by CobB during the course of reaction. Our experiments revealed also that CobB likewise becomes acetylated by acetyl phosphate in a system consisting of purified proteins (Fig. 1D). Acetylated lysine residues of the sirtuin were subsequently identified by MS and are presented in Supplementary Table 2. It remains unknown if CobB acetylation is of physiological importance. However, neither Prs nor CobB acetylation had any impact on their interaction compared to the wt and non-acetylated protein (Fig. 1E). This suggests that acetylated lysine residues of Prs do not take part in its binding to the CobB deacetylase. We further probed this by repeating the pull downs with purified Prs variants where the acetylable lysines K182, K194 and K231 were substituted by alanines. All three variants were still able to interact with CobB (Fig. 1F).

**Table 1.**
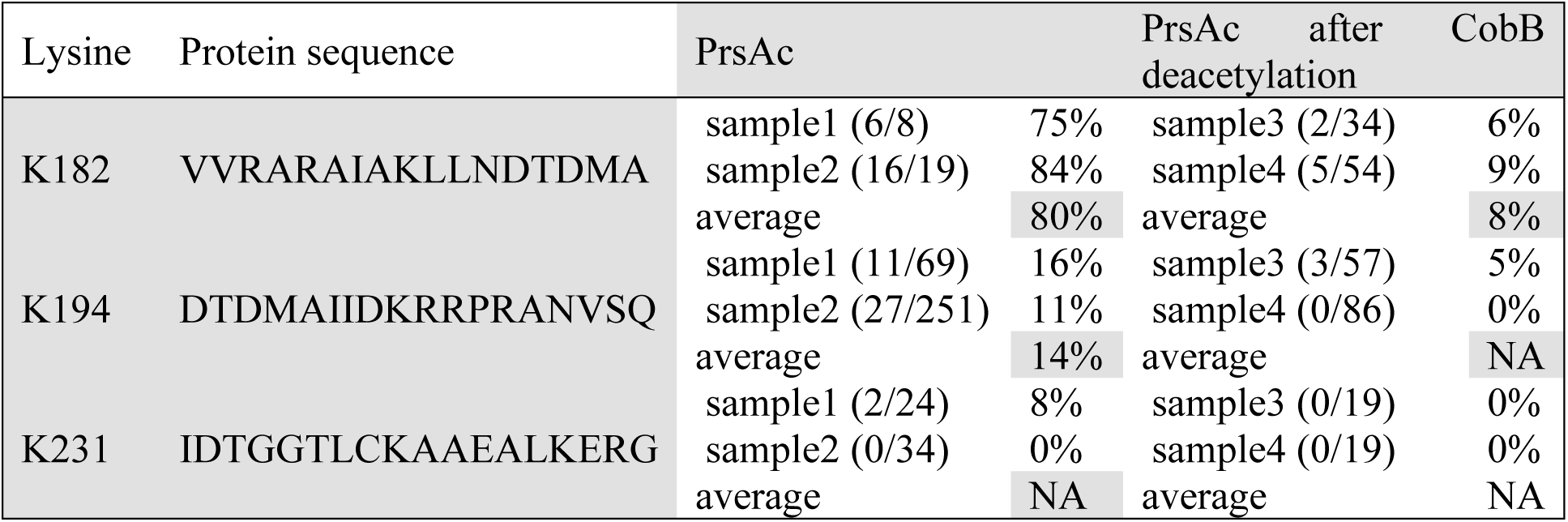
Acetylated lysines in Prs and their deacetylation by CobB. Lysine residues K182, K194, K231 are acetylated *in-vitro* by Ac-P and deacetylated by CobB deacetylase. Samples of *in vitro* acetylated Prs and subsequently deacetylated by CobB (Fig. 1D) were excised from 2 independent Coomassie stained SDS-Polyacrylamide gels and analyzed with Mass Spectrometry. All identified peptides carrying lysines of interest were summed up (second value in brackets) and acetylated lysines were counted (first value in brackets). The acetylation percentage was calculated and average values are shown in the table. A PrsAc protein sample aliquot was additionally analyzed with mass spectrometry where 108, 82 and 73 peptides with acetylated lysines K182, K194 and K231 respectively were identified suggesting those lysines were acetylated *in-vitro*.

Corroborating those results, we found that CobB was pulled-down by Prs in *ΔpatZ* and *Δpta* strains (Supplementary Table 1). The former strain is devoid of protein lysine acetyltranferase (3), the latter lacks phosphate acetylase (37) which synthesizes acetyl phosphate. The two strains are characterized by decreased level of enzymatic and non-enzymatic acetylation, respectively (4, 38).

In summary, our results have shown that Prs and CobB form a stable complex, whereas their interaction does not rely on Prs acetylation, nor is it affected by the acetylation state of both proteins *in vitro* or in cell extracts.

### Prs stimulates the deacetylase activity of CobB and partially protects it from the inhibitory effect of nicotinamide

Since CobB forms a stable complex with Prs, and its assembly is not dependent on Prs acetylation, we hypothesized that this interaction may have regulatory function for CobB. First, we investigated the effect of Prs-CobB assembly on CobB-mediated removal of acetyl groups from modified lysines. To elucidate this, we measured deacetylation rate of an artificial fluorogenic substrate for Zn2+ and NAD+ dependent deacetylases – MAL (BOC-Ac-Lys-AMC)(39). In the presence of Prs, CobB activity in deacetylating MAL was stimulated by about 50% (Fig. 2A). The degree of stimulation was dependent on Prs concentration and reached maximum at a 1:1 molar ratio of CobB to Prs (CobB monomer : Prs hexamer) (Fig. 2B). A further increase of Prs concentration had no effect on CobB activity, suggesting that Prs hexamers stimulate a single catalytic center of CobB. As expected, CobB deacetylase activity was dependent on NAD+ concentration and Prs enhanced it in a wide range of NAD+ concentrations, both lower and higher than the physiological ones (40) (Fig. 2A). Neither Prs substrates (ribose 5-phosphate, ATP) nor the products of its catalytic activity (PRPP, AMP) and allosteric inhibitor ADP, had significant impact on its binding to CobB or ameliorated deacetylation of MAL substrate (Supplementary Fig. 2).

**Figure 2.**
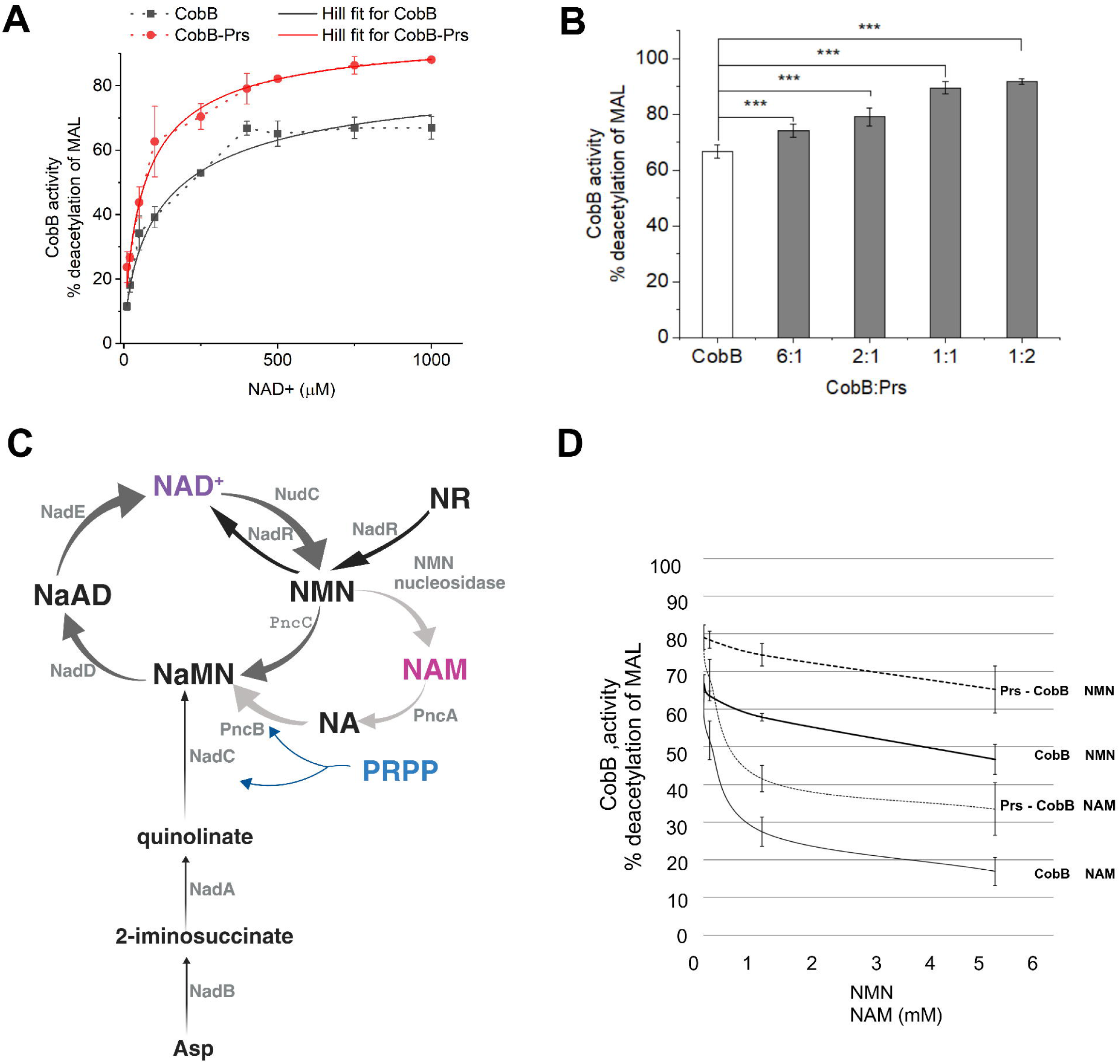
**(A) Prs stimulates CobB activity in a wide range of NAD+ concentrations. (B) Stimulation dependence on CobB:Prs (monomer:hexamer) ratio.** Deacetylation of MAL substrate (8 nmol) by CobB (320 pmol of monomer) in the presence of Prs (300 pmol of hexamer) and indicated amounts of NAD+ was performed for 1h. Fluorescent substrate was extracted with ethyl acetate and fluorescence was measured at 330/390 nm in a plate reader. Error bars represent SD between 3 independent experiments. Statistical significance was marked: *** - p-value ≤ 0.01 **(C) NAD+ producing and consuming pathways in *E. coli*.** NAD+ de novo synthesis from aspartate (thin black arrows) and salvage pathways in *E. coli*: I (represented jointly by dark and light gray arrows), II (shown by dark gray arrows), IV (from nicotinamide riboside, depicted by thick black arrows). Names of the enzymes engaged in NAD+ metabolism were provided next to the arrows depicting respective reactions. Prs catalyzes PRPP (blue) biosynthesis which serves as a substrate for NAD+ formation de-novo from aspartic acid and in NAD+ salvage pathways. NAD+ is essential for CobB mediated protein deacetylation. NAM is a byproduct of this reaction (CobB cofactor NAD+ and byproduct NAM are highlighted by different shades of purple). NAD - nicotinamide adenine dinucleotide, NMN – nicotinamide mononucleotide; NAM – nicotinamide, NA – nicotinic acid, NAMN - nicotinic acid mononucleotide, NaAD – nicotinate adenine dinucleotide, NR – nicotinamide ribonucleotide, R5P – ribose 5-phospahte, PRPP – phosphoribosyl pyrophosphate, Asp – aspartate **(D) Prs partially overcomes inhibition of CobB by NAM.** MAL deacetylation was performed as described above in the presence of indicated amounts of NAM and NMN and Prs (300 pmol of hexamer). Mean values of 3 independent experiments were presented. Error bars show SD between the repeats.

Prs produces PRPP, one of the key metabolites for the synthesis of NAD+(41) (Fig. 2C) which is a substrate of CobB in deacetylation. Therefore, a possible role of their interaction could be to orchestrate the sirtuin activity with NAD+ metabolism. CobB, as other sirtuins, was shown to be sensitive to feedback inhibition by deacetylation reaction byproduct – nicotinamide (NAM), with IC_50_ value estimated at approximately 52 μM(29) (Fig. 3C). NAM is also one of the metabolites of the *E. coli* NAD+ salvage pathway I, whereas its intracellular level has been shown to fluctuate dependent on bacterial growth conditions (29). This makes NAM a likely candidate for a regulator of CobB activity *in vivo*. Therefore, we tested how interaction with Prs affects NAM-mediated inhibition of CobB deacetylase activity. In the presence of Prs, deacetylation of MAL substrate by CobB was more effective even at very high NAM concentrations, indicating that Prs partially protects CobB from inhibition by NAM (Fig. 2D). We also confirmed using pull-down assay that CobB forms a complex with Prs in the presence of NAM (Supplementary Fig. 2).

**Figure 3.**
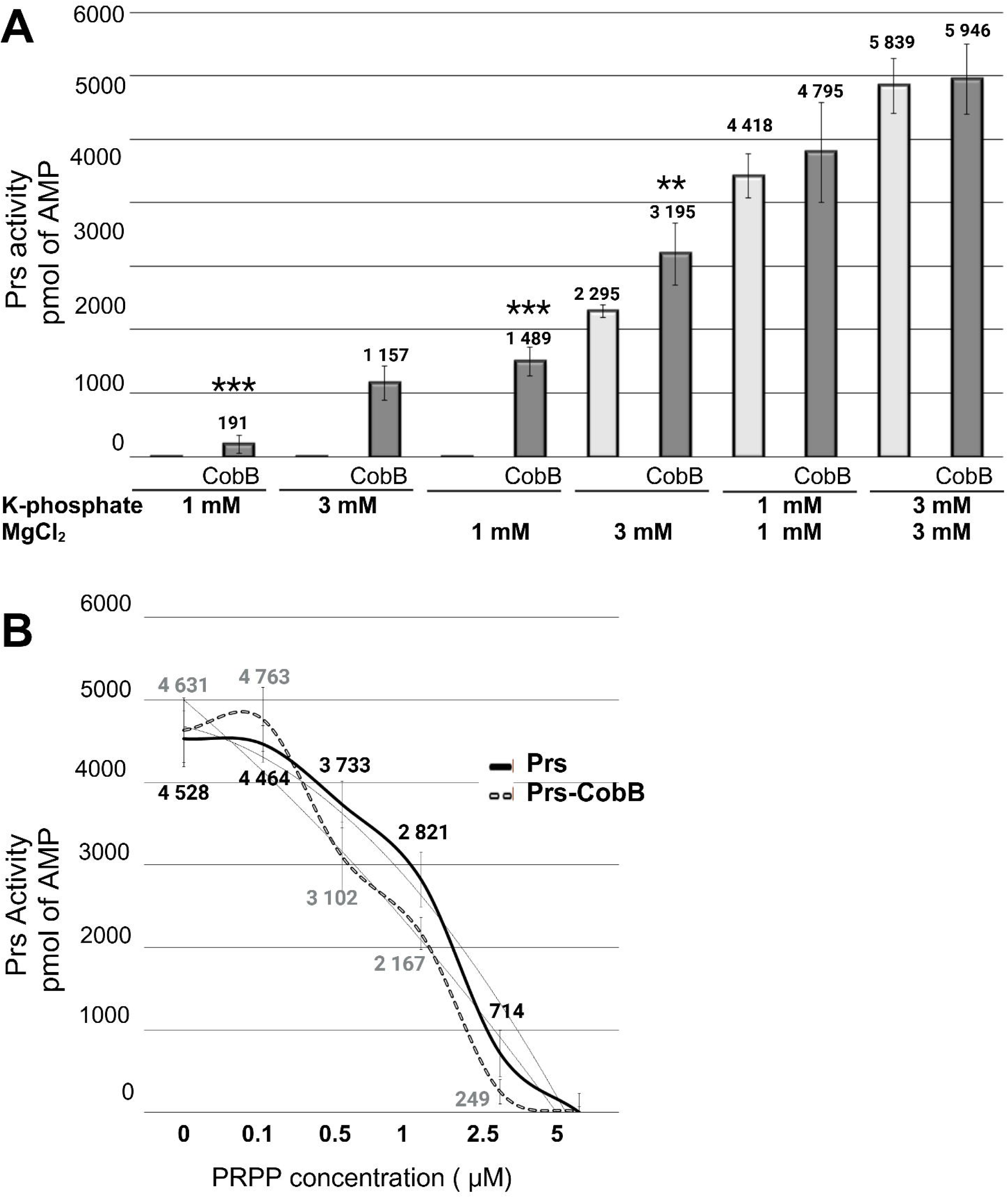
CobB-Prs interaction influences PRPP synthasès activity in suboptimal conditions. **(A) CobB stimulates Prs activity *in vitro* under conditions of low phosphate or magnesium ion concentration.** Prs activity was assessed by measuring AMP formation from ribose 5-phospate (60 μM) and ATP (60 μM). Reaction was performed in the presence of CobB (Prs hexamer : CobB monomer ratio) in 100 µl reaction buffer (50 mM Tris pH 8.0, 100 mM KCl, 0.5 mM DTT, 0.1 mg mL^-1^ BSA) with variable concentration of Mg2+ and PO4-indicated in the figure. After termination of Prs reaction, the ATP luminescence was measured using plate reader as described above. Mean values of 3 independent experiments were presented. Error bars show SD between the repeats. Statistical significance was marked for comparison of cognate samples containing either the CobB-Prs complex or Prs alone: *** - p-value ≤ 0.01; ** - p-value ≤ 0.05 **(B) CobB effect on Prs feedback inhibition by PRPP**. Prs activity was measured in 100 µl reaction buffer (50 mM Tris pH 8.0, 100 mM KCl, 1 mM MgCl_2_, 0.5 mM K-phosphate pH 8.0, 0.5 mM DTT, 0.1 mg mL^-1^ BSA) as described above in the presence of indicated concentrations of PRPP. All measurements were taken in at least 3 independent repeats. Error bars show SD between the repeats.

Moreover, it has been shown previously that several metabolites of the NAD salvage pathways, like nicotinamide mononucleotide (NMN), can act as weak inhibitors of Sir2 family deacetylases *in vitro*(38). Corroborating those results, we observed that NMN, as well as nicotinic acid adenine dinucleotide (NaAD), NADH and NADP negatively influence CobB activity at high concentrations. As in the case of NAM, Prs increased CobB activity in the presence of those metabolites (Supplementary Fig. 3).

Those results suggest that Prs-CobB interaction influences catalytic activity of CobB by increasing the deacetylation reaction rate. Formation of a complex with Prs also partially protects CobB from inhibition by NAM and other NAD+ metabolites, further suggesting that the interaction exerts an impact on the catalytic center of CobB.

### CobB stimulates Prs activity *in vitro* under suboptimal conditions

The results presented so far strongly suggest that formation of the Prs-CobB complex impacts the rate of acetyl group removal from AcK residues. We next asked, whether assembly with CobB reciprocally influences Prs activity. Prs synthesizes PRPP from ribose 5-phosphate and ATP, producing AMP as a byproduct (41). It utilizes magnesium and phosphate ions as a cofactor and allosteric activator, respectively (42). To assess Prs catalytic activity, we monitored AMP formation using a luciferase-based assay that produces a luminescent signal proportional to AMP present in the samples.

Results of this assay showed that addition of CobB alone, CobB together with its substrate NAD+ or product NAM had little impact on Prs activity, when sufficient amount of magnesium and phosphate ions was provided (Supplementary Fig. 4). Conversely, when assays were carried out under conditions of low magnesium or phosphate concentration, CobB significantly stimulated Prs activity (Fig. 3A). This suggests that under suboptimal conditions for Prs function, assembly with CobB may safeguard PRPP synthesis by Prs. The intracellular magnesium level has been previously implicated as one of the factors regulating protein acetylation in *E. coli* cells. Namely, protein acetylation was higher in cells grown in magnesium-limited media(43). This effect could be due to insufficient CobB stimulation by Prs, making magnesium ion concentration a regulator of the Prs-CobB complex activity. However, at least *in vitro*, magnesium ions concentration had no impact on the enhancement of CobB-mediated deacetylation rate by Prs (Supplementary Fig. 5). Interaction with CobB also slightly enhanced feedback inhibition of Prs activity by PRPP (Fig. 3B). Together, those results show that, at least under certain conditions, formation of the complex with CobB may also influence Prs function.

### Acetylation of Prs is physiologically important and impacts global protein acetylation

Acetylated lysine residues of Prs do not take part in the interaction with CobB, however acetylation may have an impact on the protein catalytic activity, stability, or interaction with other cellular components. Therefore, acetylation/deacetylation of Prs may still play an important role in the Prs-CobB interplay. Consequently, using non-enzymatically acetylated, purified Prs we showed that acetylation (mostly at K182 under those conditions), significantly, although not profoundly, decreases PRPP formation by Prs (Fig. 4A). Reciprocally, CobB deacetylates Prs *in vitro* and reverses that effect (Fig. 4B).

**Figure 4.**
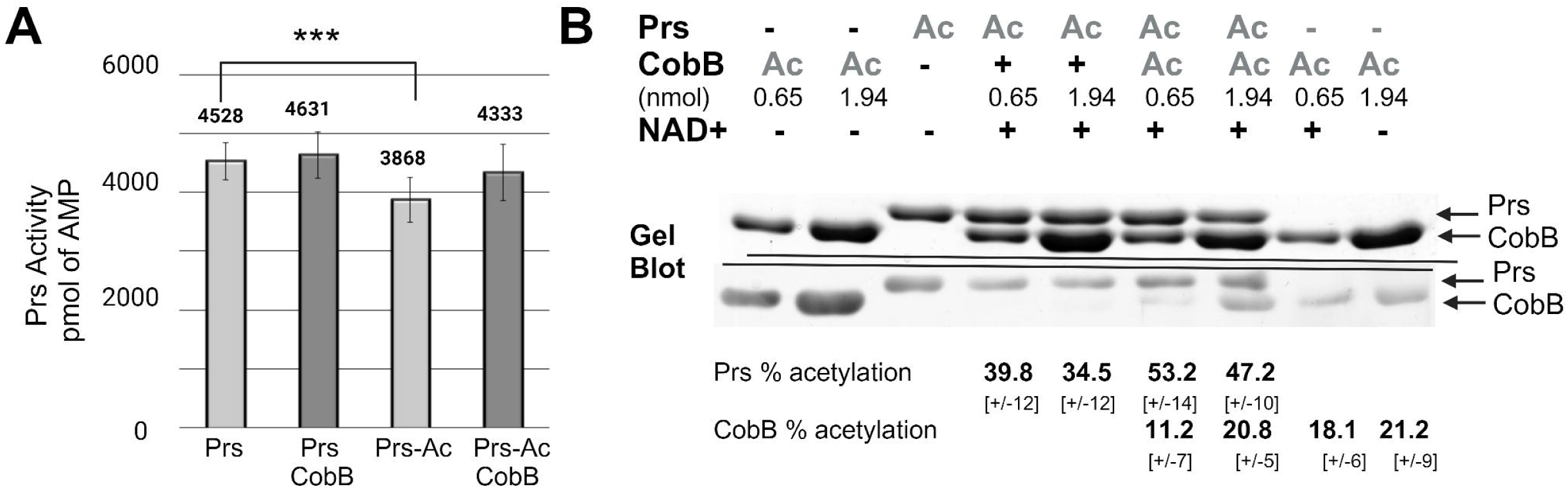
*In vitro* activity of the acetylated Prs and its deacetylation by CobB. **(A) Acetylation effect on activity of Prs.** Acetylated PrsAc exhibits slightly but significantly lower activity in comparison to non-acetylated Prs. Presence of CobB does not significantly influence the PRPP synthase activity. Prs activity (0.7 pmol) was assessed by measuring AMP formation from ribose 5-phospate (60 μM) and ATP (60 μM) in presence of CobB (4 pmol) in 100 µl reaction buffer (50 mM Tris pH 8.0, 100 mM KCl, 1 mM MgCl_2_, 0.5 mM K-phosphate pH 8.0, 0.5 mM DTT, 0.1 mg mL^-1^ BSA) After termination of Prs reaction, AMP was converted to ADP and the residual ATP was removed. Next, ATP was produced from ADP and utilized in luciferase reaction. Reactions were performed in 96-well plates and luminescence was measured using plate reader. Mean values of 3 independent experiments were presented. Error bars show SD between the repeats. Statistical significance was marked: *** - p-value ≤ 0.01

To test the role of acetylation of surface exposed K182 and K231, we introduced mutations in the chromosomal copy of the *prs* gene, resulting in changes of the lysine residues in the resulting Prs variant to alanine (A), arginine (R) or glutamine (Q). Substitution with arginine mimics lysine at permanently non-acetylated state, whereas glutamine mimics AcK residue. Using those mutants, we found that the exchange of acetylable Prs lysines to alanine or amino-acids mimicking changes to acetylation status, result in profound alterations in the global proteome acetylation level, as measured by Western blot and anti-N(ε)-acetyl lysine antibody. In those experiments, cells that chromosomally express *prs* variants K182A and K231A showed higher protein acetylation level than their wild-type counterpart upon entry to stationary phase (Fig. 5A) when protein acetylation increases (4, 38). Since Prs produces a precursor for NAD+ synthesis, changes in the enzyme’s activity could affect intracellular NAD+ concentration, ultimately leading to differences in CobB-mediated protein deacetylation rate. Therefore, product synthesis by both Prs variants was measured *in vitro*, showing that Prs K231A had a similar activity as wild-type Prs whereas the activity of Prs K182A was slightly decreased (Supplementary figure 6A). The latter result was also consistent with a slower growth rate of the cognate strain. Moreover, both variants stimulated CobB-mediated deacetylation of MAL substrate *in vitro* (Supplementary figure 6B). Mass spectrometry analysis of protein gel bands producing differential signals when probed with anti-N(ε)-acetyl lysine antibody revealed Lpd, Pgk and also GadA-GadB as their components (Supplementary table 3). All those proteins have been previously reported to be acetylated *in vivo*, whereas their acetylation level was affected in the *ΔcobB* mutant (17, 44). Thus, our data corroborate previous findings and suggests that although acetylation of Prs K182 or K231 does not influence Prs-CobB interaction (Fig. 1EF), it may affect proteome acetylation. The influence may be exerted indirectly, through metabolic changes occurring in the *prs* mutants, or directly, through the effect of *in vivo* Prs acetylation state on the Prs-CobB complex formation or deacetylase activity in the complex. Western blot results obtained with double mutants *prs* (K182A or K231A) *ΔcobB* (Fig. 5A) demonstrated that protein acetylation profile of the double mutant, upon cell entry to the stationary phase, was similar to that of the single *ΔcobB* mutant, particularly for the strain *prs* (K231A) *ΔcobB*.

**Figure 5.**
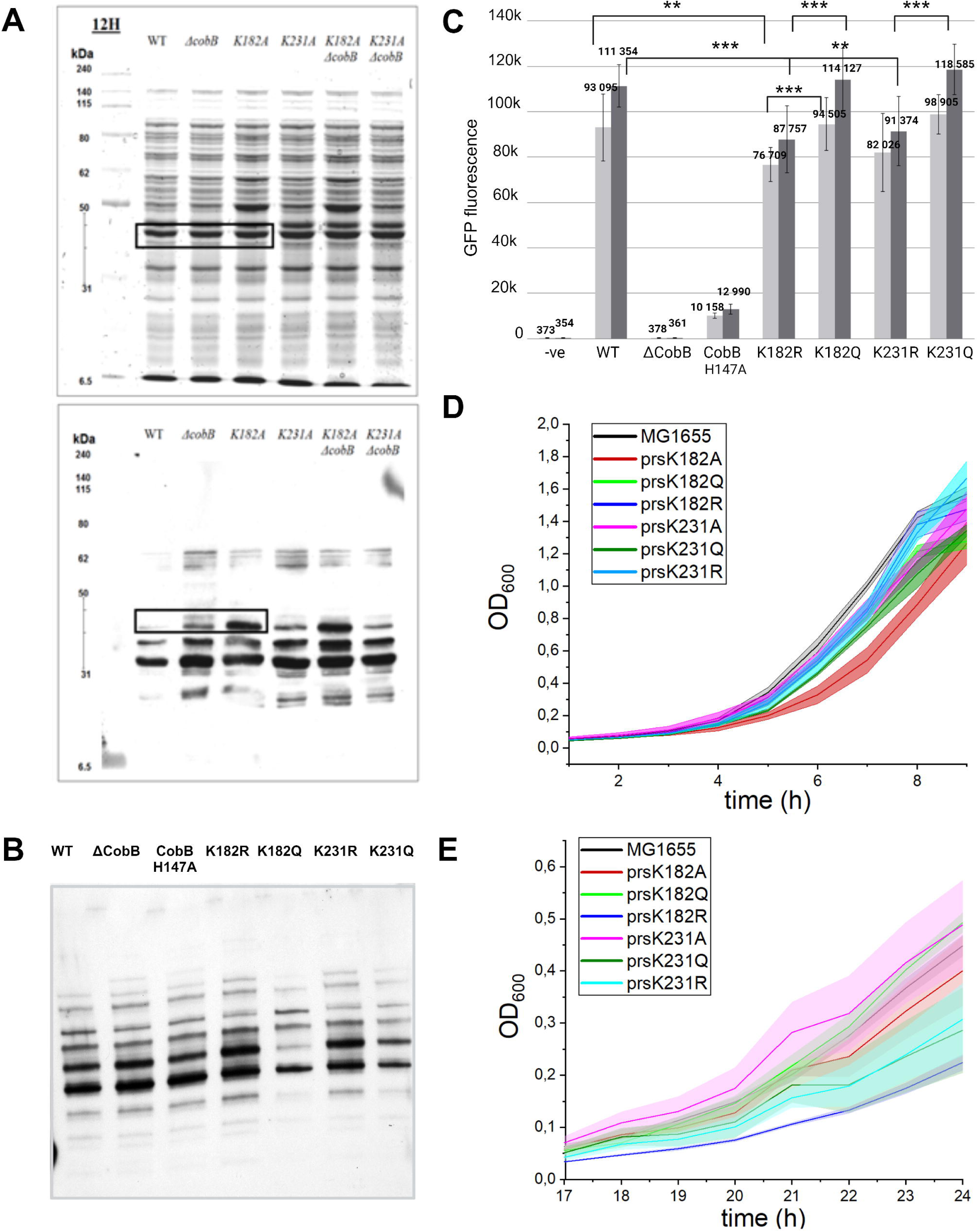
The effect of Prs K182 and K231 acetylation on global protein acetylation and *E. coli* growth. The effect of Prs variants K182A and K321A **(A)** or K182R/Q and K231R/Q **(B)** on global protein acetylation. Protein acetylation was assessed in MG1655 strain and its *ΔcobB* derivative, producing wild-type Prs or its K182A and K231A variant from the *prs* gene in its native chromosomal position. Overnight cultures of the strains were diluted in TP7.3 medium with 0.2% glucose (A) or M9 medium with 0.2% acetate (B) and samples were collected after 12 hrs of cell growth. After preparation of whole cell lysates, 6 μg of proteins from each of them was separated in 10% SDS-PAGE gel, followed by Western blot with anti-acetyl lysine antibodies. **(C)** Strains producing acetylablative Prs variants K182R and K1231R demonstrated decreased CobB activity *in-vivo*. CobB activity in prs and cobB mutant strains was measured by expression of a fluorescent probe from pULtra-AcKRS-tRNA^Pyl^-EGFP(K85TAG) plasmid allowing fluorescent EGFP production, activated upon deacetylation. The cultures were grown in LB medium supplemented with acetyl-K and appropriate antibiotics, induced at OD_600_=0.5 with 1mM Isopropyl β-D-1-thiogalactopyranoside (IPTG) and collected after 10h (light gray bars) and 14h (dark gray bars) since the induction. Fluorescence was measured for 100 µl sample in PBS in a plate reader. Mean values of 3 independent experiments were presented. Error bars show SD between the repeats. Statistical significance was marked: *** - p-value ≤ 0.01; ** - p-value ≤ 0.05 **(D) and (E) Changes mimicking Prs acetylation/deacetylation at K182 and K231 affect *E. coli* growth on different carbon sources.** Growth curve of MG1655 wild-type strain, strains with single mutations in *prs* gene replacing acetyl lysine for arginine (acetylablative) or glutamine (acetyl mimicking), and strain with *cobB* deletion and inactive cobBH147A. Overnight cultures were diluted in M9 minimal medium supplemented with 0.2 % glucose (A) or 0.2 % sodium acetate (B) and grown with aeration at 37°C. Mean values of 4 independent experiments are shown. Standard error is depicted as an error band.

Next, we tested the effect of the changes, mimicking acetylation/deacetylation of the PRPP synthase in the respective strains, as described above, after 12h cell growth in acetate medium, when overall acetylation level is high (4, 38). In those experiments, cells that chromosomally express *prs* variants K182R and K231R showed higher protein acetylation level than their K182Q and K231Q counterpart (Fig. 5B). This could suggest that enzymatic or non-enzymatic acetylation is lower or/and that CobB deacetylase activity is higher in the mutants that mimic acetylation state (Q) than in acetylation-ablative mutants (R).

To test the possible influence of Prs lysines K182 and K231 acetylation state on CobB activity *in vivo*, we used a system based on fluorescent reporter protein with genetically-encoded AcK. The efficiency of GFP fluorescence emission in this system is deacetylation-dependent and allows to assess deacetylase activity in bacterial cells(45). Results of this experiment have shown that reporter deacetylation is higher, and comparable to wt cells, in strains producing Prs variants K182Q and K231Q than in otherwise identical strains producing Prs K182R and K231R (Fig. 5C). This result suggests that lack of Prs acetylation negatively affects CobB activity. This effect could result from altered level of metabolites that stimulate or inhibit CobB, or directly from changes to Prs-CobB interaction, for instance due to differences in Prs stability.

PRPP, produced by Prs, is a necessary metabolite for *de novo* synthesis of nucleotides, cofactors, histidine and tryptophan (41), therefore altering Prs activity may have profound effect on cellular metabolism and hence bacterial growth. To verify physiological impact of Prs K182 and K231 acetylation, we tested respective mutants’ growth under conditions relevant for protein acetylation-dependent regulation, namely during glycolysis or gluconeogenic assimilation of acetate. We measured growth of the strains in a minimal medium with glucose or acetate as carbon source (Fig. 5DE). During relatively fast growth on glucose, the strain producing Prs K182A variant displayed significantly slower growth rate than the rest of strains (Fig. 5E). This variant had also lower PRPP synthase activity when measured *in vitro* (Supplementary Fig. 6A), implying that the activity of Prs might be limiting for growth on glucose. A different situation was found during bacterial growth in minimal medium supplemented with acetate. In this case, the strain producing Prs K182R showed much slower growth than the wild-type strain and the strains encoding respective Prs AcK-mimic variants (Fig. 5E). Those results strongly suggest that lysine acetylation, particularly at position K182 of Prs is relevant for bacterial growth under different metabolic regimes. The weak impact of acetylation on Prs enzymatic activity *in vitro* implies that modification of those lysines may regulate also other Prs function than that of PRPP synthase.

## Discussion

In this work we show evidence that the CobB deacetylase and the PRPP synthase form a complex in *E. coli* cells. (Supplementary Table 1, Fig. 1B) Within this complex, CobB deacetylation rate is stimulated (Fig. 2AB). Reciprocally, acetyl groups can be removed from Prs lysine residues K182, K231 and K194 by CobB (Fig. 4B). In addition, our results strongly suggest, that Prs lysines (particularly K182) and their acetylation status, play an important physiological role in *E. coli* cells. We observed differences both in global protein acetylation level and growth on diverse carbon substrates in the presence of Prs variants mimicking acetylation/deacetylation state (Fig. 5A-E). Prs acetylation had modest influence on protein’s activity *in vitro* (Fig. 4A), but *in vivo* it may affect protein stability or interaction with cellular components other than CobB, exerting more pronounced effect on PRPP synthesis. Prs acetylation state reciprocally influenced acetylation level of numerous proteins, suggesting that the underlying cause is likely partially connected to altered metabolism of acetyl group sources, like acetyl-CoA or Ac-P. However, our results also suggest that acetylation of Prs impacts CobB activity (Fig. 5C). This Prs modification likely increases during growth on acetate due to higher expression of PatZ acetylase upon relieving catabolite repression(22). In support of this notion, acetylation of Prs decreases in strains devoid of PatZ activity (17).

Structural characterization of the CobB-Prs complex will be required to identify the interaction surface and wholly uncover the complex’s role in proteome acetylation and cell physiology. Our attempts to model the interaction *in silico* and design protein changes alleviating the interaction have been unsuccessful so far.

In the experiments described in this work we have not found conditions that would break the Prs-CobB complex or restrain its formation. We have also not identified factors that would disable Prs-mediated stimulation of CobB activity *in vitro* and could therefore serve as regulators of the interaction. The known Prs inhibitors - ADP or PRPP had little effect on the cooperation between Prs and CobB (Supplementary Fig. 1). However, *prs* is regulated at transcriptional level and activated by low purine and pyrimidine nucleotides concentration (46, 47). Hence, the number of Prs-CobB complexes formed in the cell and the overall CobB deacetylation rate would depend on the demand to synthesize new purine nucleotides, including ATP precursors, that are intertwined with NAD+ synthesis. In line with such mechanism, several purine and pyrimidine synthesis enzymes, like PurA and Adk (4, 44) GuaB and PyrG(8) were found among proteins undergoing acetylation/deacetylation in *E. coli* cells. The effect of modification on those enzymes has not been determined so far, but acetylation has often an inhibitory effect. Thus, transcriptional activation of *prs* could ultimately lead to more effective protein deacetylation by CobB and, among other effects, to an increase in the activity of purine synthesis pathway. Changing levels of Prs acetylation under different metabolic regimes may also influence AcK level in a CobB-dependent manner.

In summary, the crosstalk between Prs and CobB allows *E. coli* cells to integrate various cues and adjust protein acetylation level to metabolism and NAD+ demand.

## Experimental procedures

### Materials, reagents and strains

Primers used in this study were synthesized by Sigma/Merck. A list of primers, vectors and strains is available in Supplementary table 3. Polymerases and enzymes used for cloning were purchased from Thermo Scientific or New England Biolabs. Reagents used for buffers were purchased from either Carl Roth or Bioshop Life Science. NAD^+^ salvage pathway substrates were purchased from Sigma/Merck except for nicotinamide (NAM) which was purchased from Bioshop Life Science.

Mutagenesis was performed with lambda Red recombineering as described earlier (48). Strains with single mutations replacing acetylated lysine for arginine or glutamine in *prs* gene: MG1655:*prs*K182R, MG1655:*prs*K182Q, MG1655:*prs*K231R, MG1655:*prs*K231Q; and strain with *cobB* deletion MG1655:Δ*cobB;* were constructed by recombineering in MG1655 wild-type strain (48). Strain with single mutation in active site of CobB replacing histidine for alanine: MG1655:*cobB*H147A was constructed by recombineering in MG1655:Δ*cobB* strain by introducing the mutated *cobB* allele into its primary locus

### Cloning, expression and purification of recombinant proteins

His-Prs and catalytically inactive His-PrsK194A were expressed from pET28a-His-*prs* and pET28a-His-*prs*K194A vectors in *E. coli* Rosetta (DE3) as we described earlier (49). Additional point mutations replacing acetylated lysines K182 and K231 for alanines were introduced into pET28a-His-*prs* vector with phosphorylated primers (49). *E. coli* Rosetta (DE3) was transformed with pET28a-His-*prs*K231A, grown to OD_600_ between 0.8 and 1.0 and induced with 100 µM Isopropyl β-D-1-thiogalactopyranoside (IPTG) at 37°C for 5 h in 2xYT medium (BioShop). *E. coli* BL21-DE3-pLysE was transformed with pET28A-His-*prs*K182A and induced at OD_600_ between 0.8 and 1.0 with 1 mM IPTG at 37°C for 3 h in 2xYT medium. The *cobB* sequence, amplified on *E. coli* MG1655 genomic DNA was N-terminally cloned with RF-cloning (50) into modified pET28a-TEV vector (49). The *E. coli* Rosetta (DE3) was transformed with vector pET28a-His-TEV-*cobB*, grown in Terrific Broth (BioShop) to OD_600_ between 0.8 and 1.0 and induced with 200 µM IPTG at 37°C for 5 h.

His-Prs protein and its variants were purified at 20°C, as described earlier (49) with buffer A (50 mM potassium phosphate pH 7.5, 10 % glycerol, 500 mM NaCl, 20 mM imidazole pH 7.8); buffer B (50 mM potassium phosphate pH 7.5, 10 % glycerol, 500 mM NaCl, 300 mM imidazole pH 7.8, 0.5 mM tris(2-carboxyethyl)phosphine (TCEP)) and dialysis SEC1 buffer (50 mM potassium phosphate pH 8.2, 10 % glycerol, 500 mM NaCl).

Non-acetylated CobB was purified on His-Trap columns (GE healthcare) as described above however, the lysate was loaded on the column and protein eluted on ice. Further, fractions containing protein were combined and diluted 1:1 with buffer A-0 (50 mM potassium phosphate pH 8.2, 10 % glycerol) and supplemented with Dithiothreitol (DTT) and ethylenediaminetetraacetic acid (EDTA) to a final concentration 1 mM and 0.5 mM respectively. Aliquots of 20 ml were made. TEV protease (0.2 mg) purified in-house(49) was added and incubated at 24 °C for 3 h with no shaking or rotation. After incubation, NaCl concentration was readjusted to 500 mM by spiking with 5M NaCl stock solution and protein solution was dialyzed overnight into SEC1 buffer at 4°C followed by protein separation from His tag on a His-Trap column (GE Healthcare), concentration with an Amicon Ultra 15 MWCO10kDa (Millipore) filter and size-exclusion chromatography at 4°C on a Superdex 200 10/300 GL gel filtration column (GE Healthcare). Purity of the proteins was evaluated on SDS-PAGE gel. Concentrated to 10-15 mg ml^-1^ purified proteins were snap-frozen in liquid nitrogen and stored at -70°C.

### Acetylation of His-Prs and CobB proteins during protein purification

His-Prs was purified on His-Trap columns (GE Healthcare) as described above and dialyzed overnight at 20 °C into 1 x acetylation buffer (100 mM Tris-HCl pH 7.5, 10 % glycerol, 150 mM NaCl) [modified from (5, 20)]. Further, it was concentrated using the Amicon Ultra 15 MWCO30kDa (Millipore) filter at 18°C to 4 mg ml^-1^ and acetylated with acetyl phosphate (lithium potassium salt, Sigma) at final concentration 20 mM at 24°C for 3 h. Following acetylation, His-Prs was concentrated at 18°C (Amicon Ultra 15 MWCO30kDa) to approximately 3 – 4 ml and further purified by size-exclusion chromatography at 18°C on Superdex 200 10/300 GL gel filtration column (GE Healthcare).

His-TEV-CobB, prior to acetylation, was purified and washed on His-Trap columns (GE Healthcare) with buffer Ac (50 mM Tris pH 7.9, 500 mM NaCl, 20 mM imidazole pH 7.8, 10 % glycerol) and eluted on ice with buffer Bc (50 mM Tris pH 7.9, 500 mM NaCl, 300 mM imidazole pH 7.8, 10 % glycerol). Fractions containing protein were combined, diluted 1:1 with buffer A-0 and supplemented with DTT (1 mM), EDTA (0.5 mM) and TEV protease (0.2 mg). Further, fractions were incubated for 3 h at 20°C as above, followed by overnight dialysis at 4°C into acetylation dialysis buffer (100 mM Tris, pH 8.2, 300 mM NaCl, 10 % glycerol). The CobB protein was separated from His tag on His-Trap column (GE Healthcare), concentrated at 4°C to 6 mg ml^-1^, diluted 1 : 1 in 0 x acetylation buffer (100 mM Tris-HCl pH 7.5, 10 % glycerol) and acetylated with acetyl phosphate at final concentration 20 mM at 24°C for 3 h. Protein solution was further dialyzed overnight into SEC1 buffer at 4°C.

Purity of proteins was evaluated on 10 % SDS-PAGE gel. Purified proteins, concentrated to 10-15 mg ml^-1^, were snap-frozen in liquid nitrogen and stored at -70°C. Acetylation was evaluated with Western blot probed with anti-acetyl lysine antibodies (1:800 dilution) (Sigma/Merck). Mass spectrometry outsourced to Mass Spectrometry Laboratory IBB PAS, Warsaw, Poland. Two independent batches of both His-PrsAc and CobBAc subjected to acetyl phosphate treatments were purified and acetylation was assessed.

### CobB pull-down on His-Prs and its variants

His-Prs or its variants (10 µg) were bound to 4 µl of Ni-NTA Magnetic Agarose Beads (Qiagen) in 50 µl interaction buffer (50 mM Tris pH 9.0, 30 mM imidazole pH 7.8, 300 mM NaCl, 0.2% Tween 20, 10 % glycerol) for 1 h on vibrating mixer at 20°C. Following triple wash with interaction buffer, 30 µg of CobB was added to His-Prs bound to beads in 50 µl fresh interaction buffer and incubated for 2 h on vibrating mixer at 20°C. Beads were washed 4 times with interaction buffer and resuspended in 20 µl 1 x SDS loading dye (31.25 mM Tris pH 6.8, 10 % glycerol, 1 % SDS, bromophenol blue). Proteins were resolved by electrophoresis in 10 % SDS-PAGE gel and visualized with Coomassie Brilliant Blue R250 staining. 4 µg of His-Prs and CobB proteins were loaded on gel separately as control. The experiments were performed in 3 independent repeats and representative gels were shown in figures.

### CobB deacetylase activity on MAL (BOC-Ac-Lys-AMC) substrate

CobB deacetylase activity was measured as described earlier (39, 51) as deacetylation of MAL substrate (Sigma/Merck), an artificial fluorogenic substrate for Zn2+ and NAD+ -dependent deacetylases. Briefly, deacetylation of 8 nmol MAL substrate by 320 pmol of CobB was performed for 1 h at 24 °C in 50 µl sirtuin buffer (50 mM Tris-HCl pH 8.5, 137 mM NaCl, 2.7 mM KCl, 1 mM MgCl_2_) [modified from (39)] in presence of NAD+ at final concentration 400 µM. The NAD^+^ titration was performed in sirtuin buffer supplemented with NAD^+^ independently at final concentrations 10 µM, 20 µM, 50 µM, 100 µM, 250 µM, 500 µM, 750 µM and 1 mM. The MgCl_2_ titration was performed in sirtuin buffer containing 400 µM NAD^+^ and MgCl_2_ was supplemented independently at final concentrations 10 µM, 20 µM, 50 µM, 100 µM, 250 µM, 500 µM, 750 µM, 1 mM, 5 mM and 10 mM. The effect of NAD^+^ salvage pathway substrates was evaluated by adding 5 µl of 50 mM β-nicotinamide mononucleotide (NMN), nicotinamide (NAM), nicotinic acid (NA), nicotinic acid mononucleotide (NaMN), nicotinic acid adenine dinucleotide (NaAD) or 10 mM NADH, NADP, NADPH to the reaction. The effect of Prs influence on CobB activity was measured in sirtuin buffer by addition of 150 pmol, 300 pmol and 600 pmol of Prs hexamer. The effect of Prs influence on CobB activity in NAD+ titration assay, MgCl_2_ titration assay and in presence of NAD+ salvage pathway substrates was measured by addition of 150 pmol of Prs hexamer. All reactions were stopped by addition of 400 µl of 1 M HCl and fluorescent substrate was extracted as an upper phase after vortexing for 30 s with 800 µl ethyl acetate and centrifugation (9 500 x g, 5 min). Ethyl acetate was evaporated under the hood at 65°C and the residue was dissolved in 600 µl acetonitrile buffer (39.6 % acetonitrile, 5 µM KH_2_PO_4_, 4.6 µM NaOH). The fluorescence (2 x 250 µl) was measured at 330/390 nm in black, flat bottom 96 well plate (51). The experiments were repeated in at least 3 independent replicates.

### CobB deacetylase activity on His-PrsAc

CobB driven deacetylation of His-PrsAc was analyzed by Western blot with anti-acetyl lysine antibodies (Sigma/Merck) and mass spectrometry. Briefly, CobB driven deacetylation of His-PrsAc was performed in sirtuin buffer (as above) for 1 h at 24°C in the presence of 400 µM NAD^+^. Deacetylation of 147 pmol Prs hexamer (30 µg of protein) was performed in 50 µl reaction with 650 pmol or 1940 pmol CobB or CobBAc (20 µg and 60 µg of protein respectively). The reaction was stopped by dilution in 4 x SDS loading dye and 5 minutes incubation at 100°C. Samples were resolved by electrophoresis in 10 % SDS-PAGE gel in duplicates with one gel visualized with Coomassie Brilliant Blue R250 staining and the other transferred to PVDF membrane, probed with anti-acetyl lysine antibodies (1:800 dilution) and visualized. Selected bands were excised from the gel and outsourced for identification of protein lysine acetylation by mass spectrometry to Mass Spectrometry Laboratory IBB PAN, Warsaw, Poland. The experiments were repeated in at least 3 independent replicas and representative gels are shown in figures.

### Measurement of Prs catalytic activity

The activity of Prs and its variants was measured as a luminescence signal from AMP formed during catalytic formation of PRPP from ribose-5-phosphate (60 µM) and ATP (60 µM) for 15 min at 37°C. The assay was carried out with His-Prs protein and its variants His-Prs-K194A, His-Prs-K182A and His-Prs-K231A, in 100 µl MgCl_2_ rich reaction buffer (50 mM Tris pH 8.0, 100 mM KCl, 13 mM MgCl_2_, 0.5 mM K-phosphate pH 8.0, 0.5 mM DTT, 0.1 mg mL^-1^ BSA) at final Prs hexamer concentration 7 nM (0.7 pmol of hexamer per reaction, 142.8 ng of protein per reaction).

The catalytic activity of His-Prs protein in the presence of CobB at various MgCl_2_ and potassium phosphate (pH 8.0) concentrations was measured in 100 µl reaction buffer base (50 mM Tris pH 8.0, 100 mM KCl, 0.5 mM DTT, 0.1 mg mL^-1^ BSA) at final Prs hexamer concentration 7 nM and CobB monomer concentration 40 nM (4 pmol of monomer per reaction, 124 ng of protein per reaction). Reactions were supplemented with 1 mM or 3mM potassium phosphate pH 8.0 and / or 1 mM or 3 mM MgCl_2_.

The activity of His-Prs in presence of various PRPP concentrations was measured in Prs reaction buffer (50 mM Tris pH 8.0, 100 mM KCl, 1 mM MgCl_2_, 0.5 mM K-phosphate pH 8.0, 0.5 mM DTT, 0.1 mg ml^-1^ BSA) at final Prs hexamer concentration 7 nM and CobB monomer concentration 40 nM.

Reactions were ceased by placing samples immediately after incubation on iced water. The concentration of AMP was measured with AMP-Glo™ Assay (Promega) according to manufacturer’s protocol. Briefly, the luminescence was measured for 10 µl subsample after adding 10 µl of AMP-Glo™ Reagent I which stopped the reaction, removed remaining ATP (1 h incubation at 24°C) and converted AMP produced into ADP, followed by conversion of ADP to ATP through luciferase reaction with the AMP-Glo™ reagent II (1 h incubation at 24°C). The luminescence was measured in white 96 well plate and the concentration of AMP in experimental samples was calculated based on AMP standard curve measured in triplicates. The catalytic activity of His-Prs protein variants was measured in two independent repeats while the activity in other conditions was measured in at least 3 independent experiments.

### Detection of acetylated proteins *in-vivo*

The acetylation level of the whole *E. coli* proteome was measured in M9 minimal medium (1 x M9 salts, 0.1 mM CaCl_2_, 2 mM MgSO_4_, 0.04 % thiamine, 25 g l^-1^ uridine) supplemented with 0.2% sodium acetate in early stationary phase (12 h). Bacterial cells (bacterial culture volume = 12/OD_600_) were harvested and the pellet was snap-frozen in liquid nitrogen. Frozen pellet was resuspended in 350 µl lysis buffer (30 mM Tris pH 6.8, 10 % glycerol, 100 mM NaCl, 0.2 mM tris(2-carboxyethyl)phosphine (TCEP), 1 x Pierce Protease inhibitors (ThermoFisher Scientific)) followed by 30 s sonication with 2 s :5 s pulsing : rest with 20 % amplitude in an ice-block. Samples were centrifuged 10 min at 16 000 x g, 4°C and protein concentration was measured with Bradford assay. Samples (7.5 µg) were resolved by electrophoresis in 10 % SDS-PAGE gel in duplicates, with first gel visualized with Coomassie brilliant blue and second transferred to PVDF membrane (80 min, 80 V), blocked in 7 % skimmed milk, probed with anti-acetyl lysine antibodies (1:800 dilution) and visualized. The representative gels and Western blots were repeated in at least 3 independent experiments.

### Measurement of CobB activity *in-vivo* with fluorescent probe

CobB activity in *prs* and *cobB* mutant strains was measured by expression of a fluorescent probe from the pULtra-AcKRS-tRNA^Pyl^-EGFP(K85TAG) plasmid described by (45). The amber (TAG) mutation in the EGFP at essential for fluorophore maturation lysine K85 allowed incorporation of acetylated lysine (AcK) present in media with orthogonal amber suppressor pyrrolysyl-tRNA synthetase mutant (AcKRS)/tRNA^Pyl^ pair specific for AcK. Deacetylation results in formation of a fluorescent EGFP while in the absence of deacetylases the EGFP mutant remains non-fluorescent. The chromosomally-encoded CobB activity was measured in the following strains: MG1655 wt, MG1655:*ΔcobB,* MG1655:*cobB*H147A, MG1655:*prs*K182R, MG1655:*prs*K182Q, MG1655:*prs*K231R and MG1655:*prs*K231Q, freshly transformed with pULtra-AcKRS-tRNA^Pyl^-EGFP(K85TAG) plasmid and plated on LB plate supplemented with 25 mg l^-1^ spectinomycin while MG1655:*ΔcobB* as a negative control was freshly plated on LB agar plate. A single colony was inoculated into 18 mL LB supplemented with 5 mM AcK (N(epsilon)-Acetyl-L-lysine, Alfa Aesar, J64139.03) and 25 mg l^-1^ spectinomycin (except for the negative control MG1655:*ΔcobB* which was incubated in LB supplemented with AcK only). The cultures were grown until OD_600_=0.5 and induced with 1mM Isopropyl β-D-1-thiogalactopyranoside (IPTG). Bacterial cells (bacterial culture volume = 1/OD_600_) were collected at 10 h and 14 h, washed twice in 500 µl of PBS, snap-frozen in liquid nitrogen and stored at -20°C. Further, pellets were thawed and resuspended in 500 µl of PBS and fluorescence was measured (excitation 450 nm; emission 510 nm) for 100 µl sample in a flat bottomed, transparent 96 well plate. The *in-vivo* activity of native CobB protein was measured in at least 3 independent repeats.

## Acknowledgements

The authors thank dr Ian Cadby (University of Bristol, UK) and Krzysztof Sitko (University of Gdansk, Poland) for assistance in some of the presented experiments, Prof. Peter G. Schultz lab (Scripps Research Institute, La Jolla, USA) for providing plasmid system for measuring *in vivo* sirtuins activity, and Prof. Grzegorz Węgrzyn (University of Gdansk, Poland) for critical reading of the manuscript. This work was supported by the National Science Center (Narodowe Centrum Nauki, Poland) [No. UMO-2014/13/B/NZ2/01139 to M.G. and UMO-2016/23/N/NZ2/02378 J.M-O.] and the University of Gdansk [539-D140-B858-21 and 533-D000-GF53-21 to B.W.; and 533-0C20-GS33-21 to A.S.]. Manuel Banzhaf was supported by a UKRI Future Leaders Fellowship [MR/V027204/1].

## Authors contribution

B.W. performed majority of the experimental work, contributed to work conceptualization and preparation of the manuscript; J.M-O. performed the MS analysis of protein complexes and reviewed the manuscript; A.S. performed a part of the biochemical assays; A.L. contributed to establishment of protein assays and reviewed the manuscript; J.H. contributed to manuscript writing and data visualization; M.B. contributed to project conceptualization and manuscript writing; M.G. contributed to project conceptualization, data analysis and manuscript writing.

