## Supplementary material for "Bacterial sirtuin CobB and PRPP synthase crosstalk in the regulation of protein acetylation in *Escherichia coli*": Supplemntary figures and tables

**Supplementary Table 1.** Results of AP-MS analysis of interacting proteins. Protein complexes were isolated using sequential peptide purification protocol[1]. Control samples consisted of proteins isolated from MG1655 strain grown in the specified medium, subjected to the same purification protocol but in the absence of the specific bait protein. MS data was analyzed using MaxQuant[2]. Intensity value (arbitrary counts) reflects summed peptide intensity of all identified peptides belonging to a given protein. Intensity is directly proportional to the amount of protein in the sample. Enrichment was calculated as a fold change of protein intensity in the analyzed samples in comparison to controls.

| Bait protein | Strain | Conditions | Bait intensity | Mean bait intensity | Prey | Prey intensity | Mean prey intensity | Prey intensity in the control sample | Mean prey intensity in the control samples | Enrichment (fold) |
|---|---|---|---|---|---|---|---|---|---|---|
| Prs | MG1655 | M9 + acetate | 1.87E+10<br>1.57E+10 | 1.72E+10 | CobB | 4.63E+09<br>6.56E+09 | 5.6E+09 | 3.83E+06<br>1.52E+07 | 9.51E+06 | 5.89E+02 |
| CobB | MG1655 | M9 + acetate | 3.59E+09<br>6,04E+09 | 4.81E+09 | Prs | 3.72E+09<br>4.58E+09 | 4.15E+09 | 1.37E+06<br>1.68E+06 | 1.52E+06 | 2.73E+03 |
| Prs | MG1655 | LB | 4.52E+10 | NA | CobB | 1.71E+10 | NA | 3.76E+06<br>2.89E+06<br>3.72E+06 | 3.46E+06 | 4.94E+03 |
| CobB | MG1655 | LB | 5.5E+09 | NA | Prs | 6.79E+09 | NA | 4.1E+07<br>2.3E+07<br>3.6E+07 | 3.33E+07 | 2.04E+02 |
| Prs | MG1655 *ΔpatZ* | LB | 1.72E+10<br>8.87E+09 | 1.3E+10 | CobB | 8.3E+09<br>7.56E+09 | 7.93E+09 | 3.76E+06<br>2.89E+06<br>3.72E+06 | 3.46E+06 | 2.29E+03 |
| Prs | MG1655 *Δpta* | LB | 1.28E+10<br>1.21E+10 | 1.24E+10 | CobB | 5.65E+09<br>5.87E+09 | 5.76E+09 | 3.76E+06<br>2.89E+06<br>3.72E+06 | 3.46E+06 | 1.66E+03 |
| CobB | MG1655 | LB | 5.5E+09 | NA | FabB | 2.9E+06 | NA | 2.38E+05 | NA | |
| CobB | MG1655 | LB | 5.5E+09 | NA | FabG | 1.5E+07 | NA | 6.15E+06<br>5.7E+05 | 3.36E+06 | 4.4E+01 |
| CobB | MG1655 | M9 + acetate | 3.59E+09<br>6,04E+09 | 4.81E+09 | FabB | 1.21E+06<br>2.71E+06 | 1.96E+06 | 0<br>0 | NA | NA |
| CobB | MG1655 | M9 + acetate | 3.59E+09<br>6,04E+09 | 4.81E+09 | FabG | 1.5E+07<br>1.6E+07 | 1.55E+07 | 7.2E+05<br>1.1E+06 | 9.1E+05 | 1.7E+01 |

1. Babu, M. *et al.* Sequential peptide affinity purification system for the systematic isolation and identification of protein complexes from Escherichia coli. *Methods Mol. Biol.* **564**, 373–400 (2009).

2. Cox, J. & Mann, M. MaxQuant enables high peptide identification rates, individualized p.p.b.-range mass accuracies and proteome-wide protein quantification. *Nat. Biotechnol.* **26**, 1367–1372 (2008).

**Supplementary Table 2**. CobB acetylated lysines.

Acetylation of CobB predicted with Pail[3] and measured with LC MS/MS. MS sample Ac3 consisted of liquid protein sample directly after acetylation with acetyl phosphate, while samples Ac1 and Ac2 consisted of protein bands excised from gel after electrophoresis (fig. 2D).

Ion score value indicates how well the observed MS/MS spectrum matches to the peptide stated (the higher the value the better match) while the expect value suggests the probability of accidental match between MS/MS spectra and the peptide (values below 0.1 are considered non-accidental)

| Acetylated Lysine | Amino acid sequence | Pail score | MS Sample | Query | Score | Expect |
|---|---|---|---|---|---|---|
| **K115** | AHLALA**K**LQDALG | 1.56 | Ac1 | 856 | 36 | 0.0033 |
| | | | Ac2 | 1435 | 26 | 0.2 |
| | | | | 5708 | 93 | 0.000000046 |
| | | | Ac3 | 2573 | 60 | 0.00007 |
| | | | | 2574 | (38) | 0.14 |
| | | | | 7765 | (89) | 0.000000097 |
| | | | | 7766 | 121 | 0.000000000067 |
| | | | | 13678 | 121 | 0.000000000058 |
| | | | | 14503 | 15 | 1 |
| **K152** | MHGELL**K**VRCSQS | 0.22 | Ac1 | 4011 | (43) | 0.0011 |
| | | | | 4014 | 24 | 0.093 |
| | | | | 4015 | 33 | 0.012 |
| | | | Ac3 | 9546 | 56 | 0.00027 |
| **K271** | FVE**K**LLKGLKAGS | 1.01 | Ac1 | 16 | (53) | 5.7e-05 |
| | | | | 17 | 53 | 5.7e-05 |
| | | | | 18 | (48) | 0.00019 |
| | | | Ac3 | 28 | (42) | 0.0041 |
| | | | | 29 | 42 | 0.0036 |
| | | | | 10392 | 98 | 1.3e-08 |
| **K274** | KLLKGL**K**AGSIA | 1.58 | Ac1 | 81 | 40 | 0.0039 |
| | | | Ac3 | 202 | 29 | 0.23 |
| **K32** | D**K**VVPEAMEKPR | | Ac1 | 1613 | (46) | 0.00029 |
| | | | | 2800 | (44) | 0.0006 |
| | | | | 2801 | (52) | 0.0001 |
| | | | | 2802 | (62) | 1.1e-05 |
| | | | | 2803 | 79 | 2.3e-07 |
| | | | | 2882 | 4 | 6.7 |
| | | | Ac2 | 3934 | 8 | 18 |
| | | | | 3935 | 7 | 19 |
| | | | | 3938 | 3 | 46 |
| | | | | 4026 | (52) | 0.0005 |
| | | | | 4027 | (62) | 5e-05 |
| | | | | 4028 | 32 | 0.05 |
| | | | Ac3 | 5328 | (80) | 8.8e-07 |
| | | | | 5329 | (75) | 3.6e-06 |
| | | | | 5330 | 20 | 0.95 |
| | | | | 5332 | (82) | 5.8e-07 |
| | | | | 5333 | (59) | 0.00013 |
| | | | | 5334 | (54) | 0.00042 |
| | | | | 5336 | 7 | 21 |
| | | | | 5338 | (70) | 1.1e-05 |
| | | | | 5339 | (68) | 1.6e-05 |
| | | | | 5340 | 5 | 27 |

| | | | | | | |
|---|---|---|---|---|---|---|
| | | | | 5341 | (52) | 0.00065 |
| | | | | 5485 | 15 | 2.5 |
| **K40** | VVPEAME**K**PR | | Ac1 | 1705 | 65 | 4.4e-06 |
| | | | | 1706 | 24 | 0.053 |
| | | | Ac2 | 1267 | (51) | 0.00048 |
| | | | | 2581 | (80) | 5.9e-07 |
| | | | | 2582 | (39) | 0.01 |
| | | | Ac3 | 1545 | (48) | 0.00082 |
| | | | | 1546 | (63) | 2.4e-05 |
| | | | | 3344 | (62) | 4.1e-05 |
| | | | | 3346 | (79) | 8.3e-07 |
| | | | | 3351 | (73) | 3e-06 |
| | | | | 3553 | (77) | 1.3e-06 |
| | | | | 3554 | (74) | 2.2e-06 |
| | | | | 5335 | 20 | 1.1 |
| | | | | 5337 | 2 | 64 |
| | | | | 5486 | (36) | 0.02 |

3. Li, M. *et al*. Prediction of Nε-acetylation on internal lysines implemented in Bayesian Discriminant Method. *Biochem Biophys Res Commun*. 350, 818-824 (2006).

**Supplementary Table 3.** MS protein identification of bands, excised from proteome acetylation gels (Fig. 4), which showed high acetylation signal on Western Blot.

| | | GadA 53.2kDa | GadB 53.2 kDa | lpdA 50.9 kDa | pgk 41.2 kDa |
|---|---|---|---|---|---|
| MG1655-1 | Protein Score | 3750 | 3711 | 70 | not found |
| | emPAI | 43.56 | 43.56 | 0.09 | n/a |
| | seq coverage | 58% | 58% | 2% | n/a |
| MG1655-2 | Protein Score | 1137 | 1080 | 3772 | 4499 |
| | emPAI | 3.55 | 3.55 | 14.54 | 89.42 |
| | seq coverage | 38% | 38% | 51% | 76% |
| MG1655:prsK182A-1 | Protein Score | 8005 | 53204 | 114 | not found |
| | emPAI | 488.8 | 574.29 | 0.09 | n/a |
| | seq coverage | 74% | 75% | 3% | n/a |
| MG1655:prsK182A-2 | Protein Score | 8523 | 8536 | 2392 | 2321 |
| | emPAI | 172.82 | 202.72 | 6.92 | 14.69 |
| | seq coverage | 74% | 74% | 53% | 42% |
| MG1655:ΔcobB | Protein Score | 1894 | 1838 | 3927 | 3699 |
| | emPAI | 4.34 | 4.34 | 22.62 | 31.59 |
| | seq coverage | 37% | 37% | 59% | 58% |

**Supplementary Table 4.**

**Primers used in the study**

| **primer name** | **description** | **sequence** |
| --- | --- | --- |
| K182A-F | phosphorylated, point mutation in pET28A-His-*prs* | TGCCCGCGCTATCGCTGCGCTGCTGAACGATACCGATATGG |
| K231A-F | phosphorylated, point mutation in pET28A-His-*prs* | TGGCGGTACGCTGTGTGCAGCTGCTGAAGCTCTGAAAGAACG |
| RF_pET28a-*cobB*-Nter-TEV_F | cloning of *cobB* gene into pET28A-TEV vector | CAGCGAGAATTTGTATTTTCAGGGTCGACAAGCTATGCTGTCGCGTCGGGGT |
| RF_pET28a-*cobB*-Nter-TEV_R | | TGGTGCTCGAGTGCGGCCGCATCAGGCAATGCTTCCCGC |
| RF-pkD13-prs_F | cloning of prs with point mutations into pkD13 vector | TGTCAAACATGAGAATTAATTCCGGGGATCCGTGCCTGATATGAAGCTTTTTGC |
| RF-pkD13-prs _R | | GGAACTTCGAACTGCAGGTCGACTTAGTGTTCGAACATGGCAGA |
| prs-change-cas_F | amplification of insert for lambda Red recombineering on pKD13-prsK-A* | GTTTTCGGCAGATTCTTTCCACCAATGGACGCATGCCTGAGGTTCTTCTCGTGCCTGATATGAAGCTTTTTG |
| prs-change-cas_R | | ATATTACAGACAAAAAAACCCGCCGCAGCGGGTCTTTGAGCCGGGTTCGAGCTGGAGCTGCTTCGAAG |
| cobB-KNO-cas_F | amplification of insert for lambda Red recombineering deletion of cobB, amplified on pKD13 | TGCGTGGTGCGGCCTTCCTACATCTAACCGATTAAACAACAGAGGTTGCTTGTAGGCTGGAGCTGCTTCG |
| cobB-KNO_cas_R | | CGCAAATTCAATTAATTGCGTCCCCTTGCAGGCCTGATAAGCGTAGTGCAGTCCATATGAATATCCTCCTTAGTTCC |

**Vectors used in this study**

| **vector name** | **orgin** | **description** |
| --- | --- | --- |
| pET28a-TEV | Walter et al. 2020[4] | modified pET28a protein expression vector with TEV cleavage site enabling cloning cleavable 6xHis proteinN-terminally or C-terminally |

| | | |
|---|---|---|
| pET28a-His-*prs* | Walter et al. 2020[4] | *prs* gene N-terminally cloned into pET28a |
| pET28a-His-*prs*K194A | Walter et al. 2020[4] | *prsK194A* gene N-terminally cloned into pET28a |
| pET28A-His-*prs*K182A | this study | *prsK182A* gene N-terminally cloned into pET28a |
| pET28a-His-*prs*K231A | this study | *prsK231A* gene N-terminally cloned into pET28a |
| pET28a-His-TEV-*cobB* | this study | cleavable C-terminally *cobB* cloned into pET28a |
| pKD13-*prs*K182A | this study | prsK182A gene cloned for FRT-KanR-FRT amplification of lambda red insert amplification |
| pKD13-prsK231A | this study | prsK231A gene cloned for FRT-KanR-FRT amplification of lambda red insert amplification |

**Strains used in this study**

| **strains** | **orgin** | **description** |
|---|---|---|
| DH5α | CGSC | non-mutagenized derivative of DH1, laboratory strain used for cloning |
| MG1655 | CGSC | *E. Coli* E12 laboratory strain |
| *E. coli* Rosetta (DE3) | Merck/Sigma | F- *ompT hsdS*B(rB- mB-) *gal dcm* (DE3) pRARE (CamR) |
| *E. coli* BL21-DE3-pLysE | Merck/Sigma | F- *ompT hsdS*B(rB- mB-) *gal dcm* (DE3) pLysE (CamR) |

| | | |
|---|---|---|
| MG1655:*prs*K182A | this study | MG1655 strain with point mutation in *prs* gene replacing lysine K182 for alanine with lambda Red recombineering with KanR removed with Flp-FRT recombination |
| MG1655:*prs*K231A | this study | MG1655 strain with point mutation in *prs* gene replacing lysine K231 for alanine with lambda Red recombineering with KanR removed with Flp-FRT recombination |
| MG1655:Δ*cobB* | this study | MG1655 strain with *cobB* deletion with lambda Red recombineering with KanR removed with Flp-FRT recombination |
| MG1655:*prs*K182A:Δ*cobBKanR* | this study | MG1655 strain with point mutation in *prs* gene replacing lysine K182 with lambda Red recombineering with KanR removed with Flp-FRT recombination for alanine and cobB replacement with lambda Red recombineering with KanR |
| MG1655:*prs*K231A:*Δc obBKanR* | this study | MG1655 strain with point mutation in *prs* gene replacing lysine K231 for alanine with lambda Red recombineering with KanR removed with Flp-FRT recombination and cobB replacement with lambda Red recombineering with KanR |

4. Walter, B. M., Szulc, A., and Glinkowska, M. K. (2020) Reliable method for high quality His-tagged and untagged E. coli phosphoribosyl phosphate synthase (Prs) purification. *Protein Expr Purif*. 10.1016/j.pep.2020.105587

```
-----------------------------------MPDMKLFAGNATPELAQRIANRLYT 25
                                      :*.* **:   **  ::  *

SLGDAAVGRFSDGEVSVQINENVRGGDIFIIQSTCAPTNDNLMELVVMVDALRRASAGRI 85
:    :  *.***: . :  **** * *::**** *:*::.::*.:::*:: ****.*:

TAVIPYFGYARQDRRVRSARVPITAKVVADFLSSVGVDRVLTVDLHAEQIQGFFDVPVDN 145
**::*****:***** *  ****::::*:.:: :*:*::** ***::*******:****

VFGSPILLEDMLQ-LNLDNPIVVSPDIGGVVRARAIAKLLNDTDMAIIDKRRPRANVSQV 204
 *.: ::.  :          : :*****.::**** **:: : ::**:****   * :::

MNIIGDVDGRDCIMVDDMIDTGGTLCQAAAALKAKGAKRVFAYATHPVFSGNAANNITES 264     R9PM74_Agarivorans_albus
MHVIGEVFNRDCILVDDMIDTGGTLCQAAEALKKRGARRVFAYATHPIFSGNAVKNLQNS 264     P59512_Buchnera_aphidicola
MNIIGDVAERDCVLVDDMIDTGSTLCNAAEALKYRGAKRVFAYTTHPIFSGNAVNNLRNS 264     A0A451DJZ1_Candidatus_Erwinia_haradaeae
MHIIGDVAGRDCVLVDDMIDTGGTLCKAAEALKERGAKRVFAYATHPIFSGNAANNLRNS 264
*::**:*  *:*.::**:***..*: :*: .**   * .*  *::* ::** *  .:

VIDEVVVCDTIPLSDEIKSLPNVRTLTLSGMLAEAIRRISNEESISAMFEH--        315
:* .:  *:  :   .     :    :: :::* :: : *:**:* :*
```

 * conserved in all 245 strains alligned

: 2 variants of amino acids appears

[  ] (blank) >2 variants of amino acids appears

Supplementary Fig. 1. *prs* sequence of 244 strains alligned against the E. coli strain K12 *prs*. Positions of conserved K182 and K194 are highlighted in blue. K231 is marked in green and alternative amino acids found in this position in 3 strains are shown in red

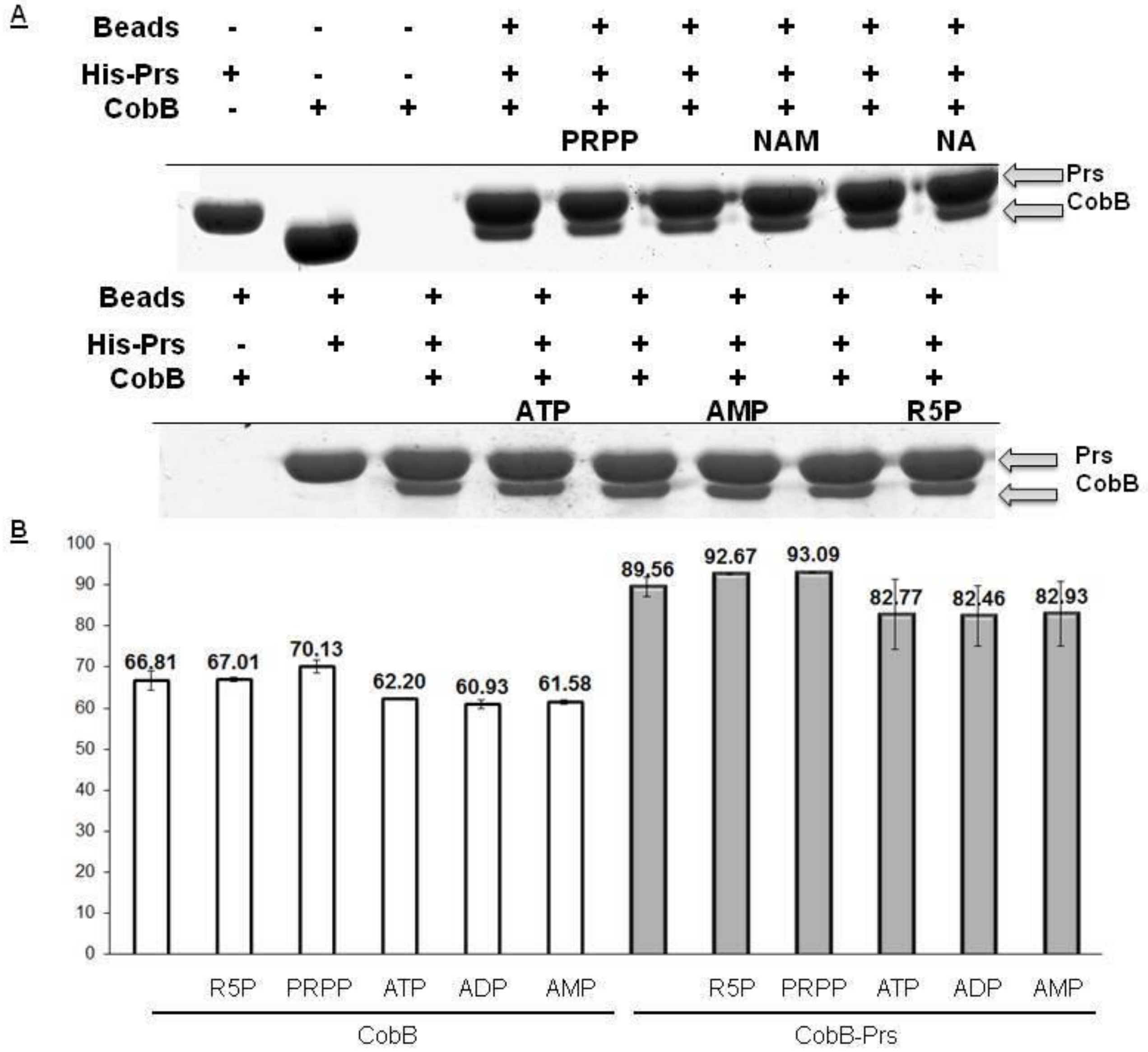


**Supplementary figure 2. Prs substrates and products of its catalitic activity do not influence CobB binding and activity.**

(A)

His6-Prs interact s with CobB in the presence of PRPP, NAM, NA, ATP, AMP and R5P substrates and products in a pull down assay.

His

6 Prs was pre bound to $Ni^{2+}$ coated magnetic beads After washing off the excess of Prs the beads were incubated with CobB Unbound protein was removed by washing and the beads were resuspended in a loading dye and separated in 10% SDS PAGE gel.

(B) R5P, PRPP, ATP, ADP and AMP do not influence CobB and CobB in complex with Prs mediated
deacetylation of MAL substrate.
Deacetylation of MAL substrate 8 nmol by CobB 320 pmol in the presence of Prs 300 pmol (of hexamer)
was performed for 1 h. Fluorescent substrate was extracted with ethyl acetate and fluorescence was measured
at 330/390 nm in a plate reader. Error bars represent SD between 3 independent experiments.

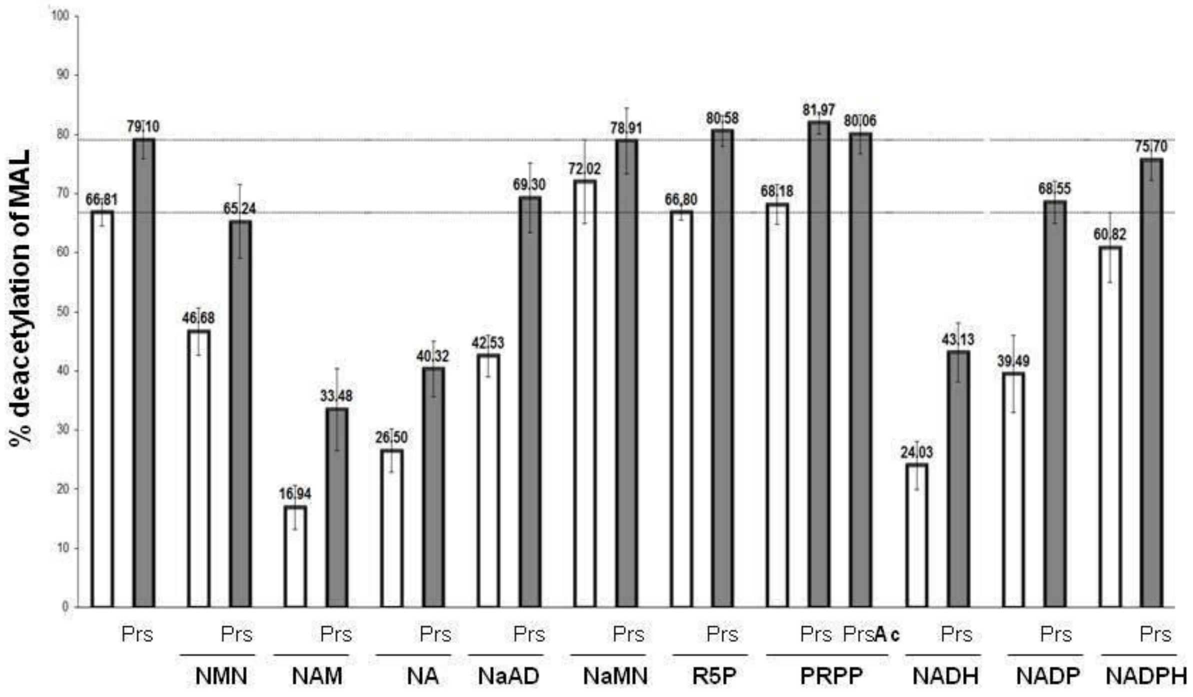


**Supplementary figure 3. CobB activity as a deacetylation of fluorogenic substrate MAL in presence of Prs and NAD+ metabolites.** NMN - nicotinamide mononucleotide, NAM – nicotinamide, NA - nicotinic acid, NaAD - nicotinate adenine dinucleotide, NaMN - nicotinic acid mononucleotide, R5P-ribose 5 phospahte, PRPP - phosphoribosyl pyrophosphate, NADH, NADP and NADPH. Deacetylation of MAL substrate 8 nmol by CobB 320 pmol in the presence of Prs 150 pmol of hexamer was performed for 1 h. Fluorescent substrate was extracted with ethyl acetate and fluorescence was measured at 330/390 nm in a plate reader. Error bars represent SD between 3 independent experiments.

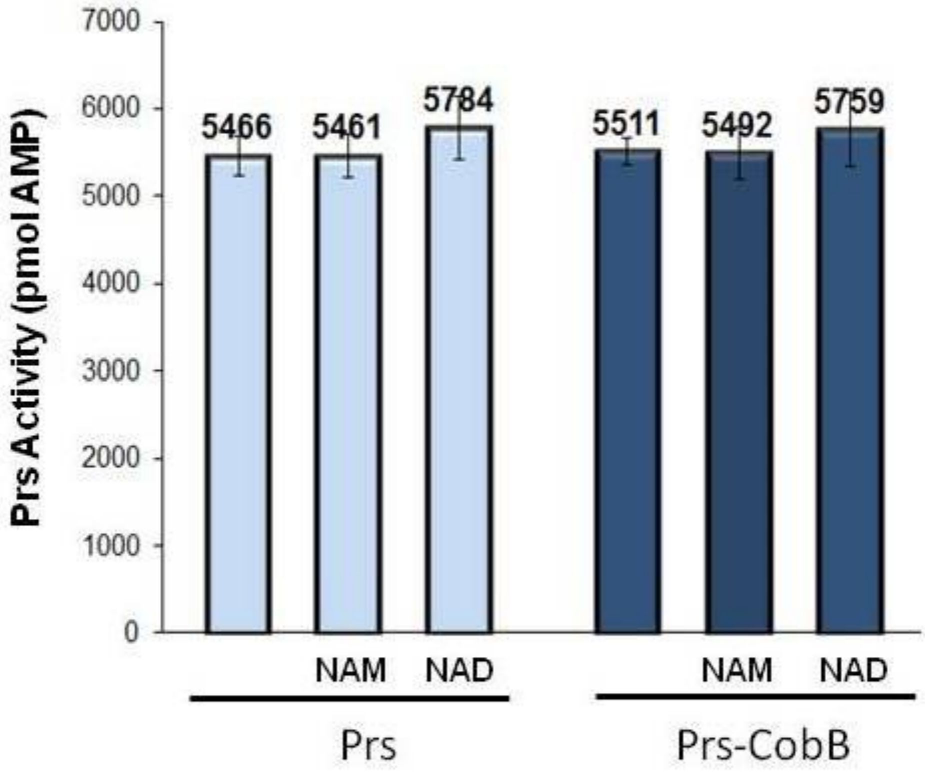


**Supplementary figure 4. Prs activity in presence of NAM and NAD**

Prs activity 25 pmol was assessed by measuring AMP formation from ribose 5-phospate 60 μM and ATP 60 μM in presence of CobB 16 pmol in 100 μl $MgCl_2$ rich reaction buffer 50 mM Tris pH 8 0 100 mM KCl 13 mM $MgCl_2$ 0.5 mM K-phosphate pH 8 0 0 5 mM DTT, 0 1 mg mL 1 BSA) After termination of Prs reaction, AMP was converted to ADP and the residual ATP was removed Next, ATP was produced from ADP and utilized in luciferase reaction Reactions were performed in 96 well plates and luminescence was measured using plate reader. Mean values of 2 independent experiments were presented. Error bars show SD between the repeats.

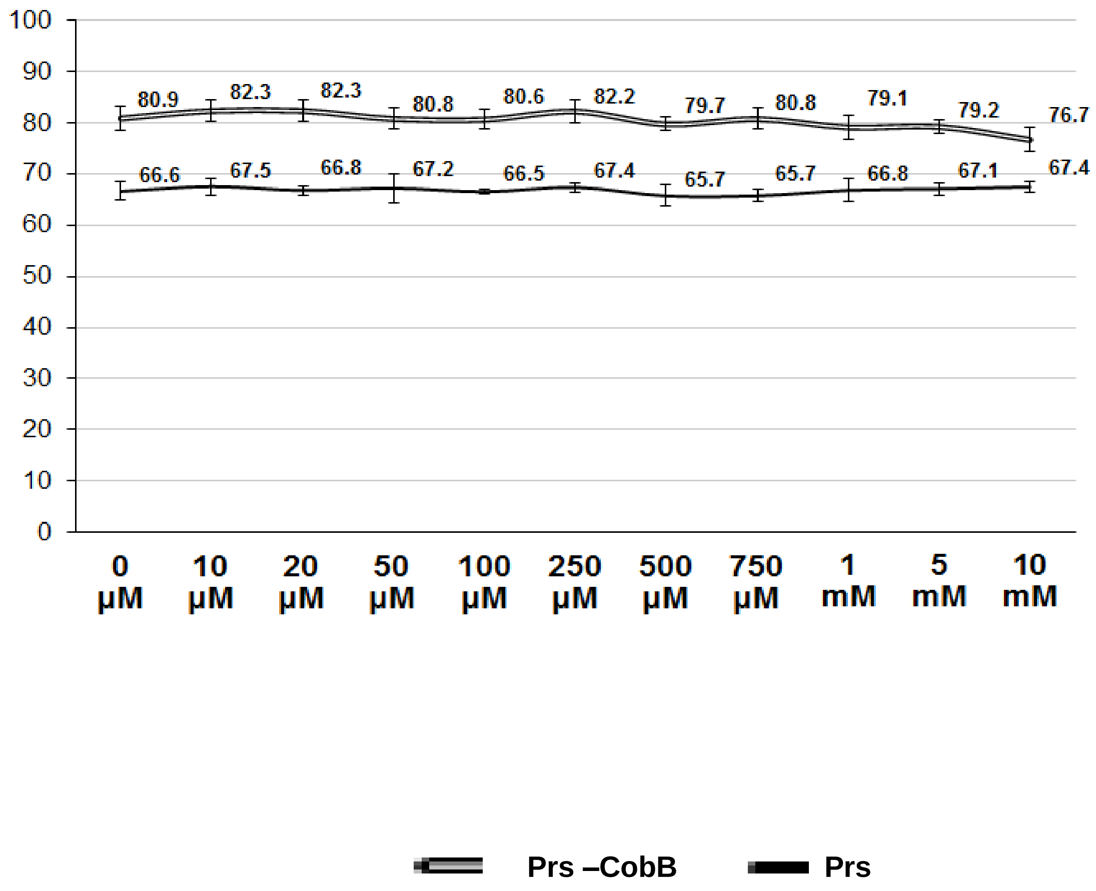


**Supplementary Figure 5. CobB activity is not affected by $MgCl_2$ concentration.**

Deacetylation of MAL substrate (8 nmol) by CobB (320 pmol) and in the presence of Prs (150 pmol of hexamer) was performed for 1h in sirtuin buffer containing 400 µM $NAD^+$ and $MgCl_2$ was supplemented independently at various concentrations. Fluorescent substrate was extracted with ethyl acetate and fluorescence was measured at 330/390 nm in a plate reader. Error bars represent SD between 3 independent experiments.

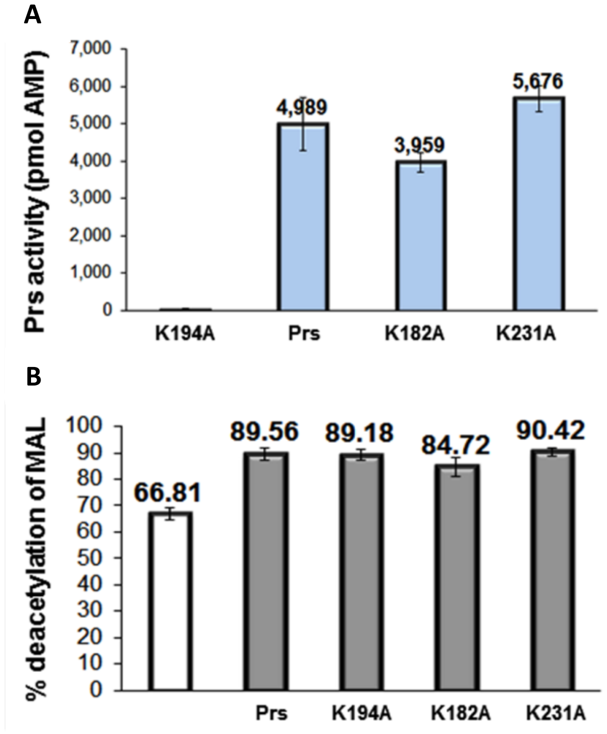


**Supplementary Figure 6. Prs variants activity *in vitro***

(A) Prs variants K182A and K231A are active in PRPP synthesis *in vitro*. Prs activity was assessed by measuring AMP formation from ribose 5-phospate (60 µM) and ATP (60 µM). After termination of Prs reaction, AMP was converted to ADP and the residual ATP was removed. Next, ATP was produced from ADP and utilized in luciferase reaction. Reactions were performed in 96-well plates and luminescence was measured using plate reader. Mean values of 2 independent experiments were presented. Error bars show SE between the repeats.
(B) Prs variants K182A, K194A and K231A stimulate CobB deacetylase activity *in vitro*. Deacetylation of MAL substrate (8 nmol) by CobB (320 pmol) in the presence and absence of Prs (6 µM as a hexamer) was performed for 1h. Fluorescent substrate was extracted with ethyl acetate and fluorescence was measured at 330/390 nm in a plate reader. Mean values of 3 independent experiments were presented. Error bars show SD between the repeats.
